# Developing a robust technique for denoising and extracting speech and breath sounds in voice recordings from asthmatic patients

**DOI:** 10.1101/2023.01.20.524994

**Authors:** Sania Fatima Sayed, Faisal I. Rezwan

## Abstract

Auscultation and processing cough, voice and breath sounds play an important role in diagnosis of several pulmonary ailments. There have been a number of studies using machine learning algorithms on such sound files to build classification and prediction algorithms. Since these studies used specialized microphones in controlled environments, it is difficult to test and deploy these algorithms in real-life settings. Recorded speech files consist of breath and wheeze sounds and it is challenging to extract from this single sound file. Hence, several audio processing and editing software are used to demarcate these sounds. The proposed technique uses a combination of a denoiser and an extraction technique to overcome these drawbacks. The developed pipeline ensures that the audio files are free of any environmental and background noises, and the audio can be recorded through any kind of microphone and environmental settings. The extraction technique further is the result of combinations of filters to output the speech and breath sounds as individual sound files, ready for processing and eliminating the need of audio editing and processing software.

## Introduction

Breath and speech demarcation and extraction from audio signals play an important role in multiple applications like speech recognition, speaker identification, and voice analysis. In recent studies, these methods have been utilized in the disease diagnosis domain. Out of many important applications, few can be seen in respiratory abnormality detection for ailments, such as bronchitis, asthma, COPD, pertussis and COVID-19 diagnosis. More than 10 databases are publicly available for such research [1]. This type of data includes cough sounds [2, 7 - 12], lung sounds [3, 5, 6], breathing sounds [3, 11, 12] and voice recordings [4]. These sounds have been recorded either by stethoscope [3, 5,6] in case of lung sounds and using specialized microphones in other cases [2, 4, 7 -11]. The Coswara, a public dataset, uses smartphone microphones to collect data openly through their web application. This was designed to overcome the drawback of implementing their COVID-19 model in a real-time environment [11]. Hence, denoising the sound files is an essential task to be able to execute the models in clinical practices where environmental and background noise is evident. Instead of recording cough, breath and voice sounds separately, in this study we have used recorded speech files from subjects with physician-diagnosed mild atopic asthma reading standard text. This proposed pipeline can be used to pre-process this data to first denoise the audio files and then separate the speech sound from the breath sounds.

## Methodology

### Dataset

The data used in this study are 10 recorded samples from non-smoking, clinically stable subjects with physician-diagnosed, and mild atopic asthma. These participants underwent a standardized inhaled methacholine challenge and after each challenge, the participants read a standardized text for 30 seconds into an external microphone.

### Denoising the sound files

These sound files had environmental and background noises like buzz sounds as the patient was speaking. For the proper filtration of sound files further into breath and speech sounds and to make the model, device independent i.e., to enable the recording of the sound files even in smartphones and in environments with noise. In this step, the denoiser utilized the Demucs Architecture proposed by Facebook AI Research [14]. This architecture uses an encoder/decoder, which is composed of convolutional encoder, a bidirectional LSTM with 2 layers, a convolutional decoder, with the encoder & decoder linked with the skip U-Net connections. The audio file was processed through this architecture to give a clean and noise free output.

### Separation of breathing and speech segments from sound files

In the vocal separation phase, the attempt was to separate the voice and the breath sounds in the denoised audio files. This technique is based on the REpeating Pattern Extraction Technique (REPET) method developed by Rafil et al. [15] with modifications which are as follows: the fast Fourier Transform (FFT) windows used overlap by ¼, rather than ½ and non-local filtering is then converted into a soft mask Wiener filter as given by Fitzgerald et al. [16]. The steps include loading the denoised audio, plotting its spectrum and Mel-Frequency Cepstral Coefficient (MFCC) to verify the difference between the original sound file and the denoised sound file. Further by using cosine similarity, similar frames are aggregated by taking per-frequency median value which are stored to be the raw filter [15]. The frames are separated by 5 seconds. Two soft mask Weiner filter is then prepared to give a foreground and a background mask. The background mask is a product of the raw filter, product of the margin given to reduce bleeding and difference between full spectrum and raw filter, and associated power. While the foreground mask is the product of the difference of the full spectrum and raw mask, margin multiplied to raw filter, and power.

*Foreground Mask = librosa*.*util*.*softmask(S_filter, margin_i * (S_full – S_filter), power = 10)*

*Background Mask = librosa*.*util*.*softmask(S_full - S_filter, margin_v * S_filter, power = 2)*

Where, S_filter is the raw filter, S_full is the full spectrum, margin_i and margin_v are the margins specified to reduce bleeding of signal, and power given for the mask.

The masks are then multiplied with the full spectrum, to give the foreground and the background signal. The foreground signal represents the filtered voice sounds, and the background signal is the remaining signal that consists of the breath sounds.

## Results

It can be observed that noise present in Figure 3a is removed in Figure 3b, it is evident from the signal that the noise present between the signals looks diminished.

**Fig 2:**
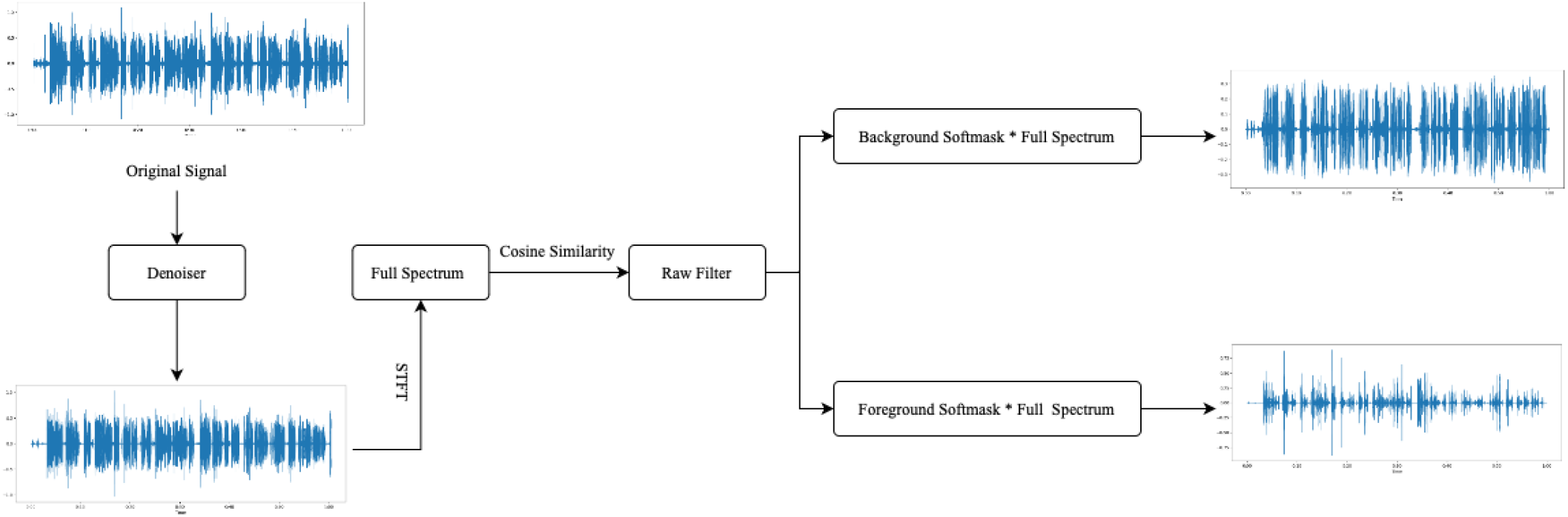
Flowchart for the entire technique. The process includes the initial original recording being denoised and the product is then being processed through STFT. The Full Spectrum is then again undergoing cosine similarity and a raw filter is created. The Raw Filter is then distinguished as Background Softmask and Foreground Softmask with their respective operations therefore multiplying with the Full Spectrum to result in the Background and Foreground signal respectively.

**Fig 3:**
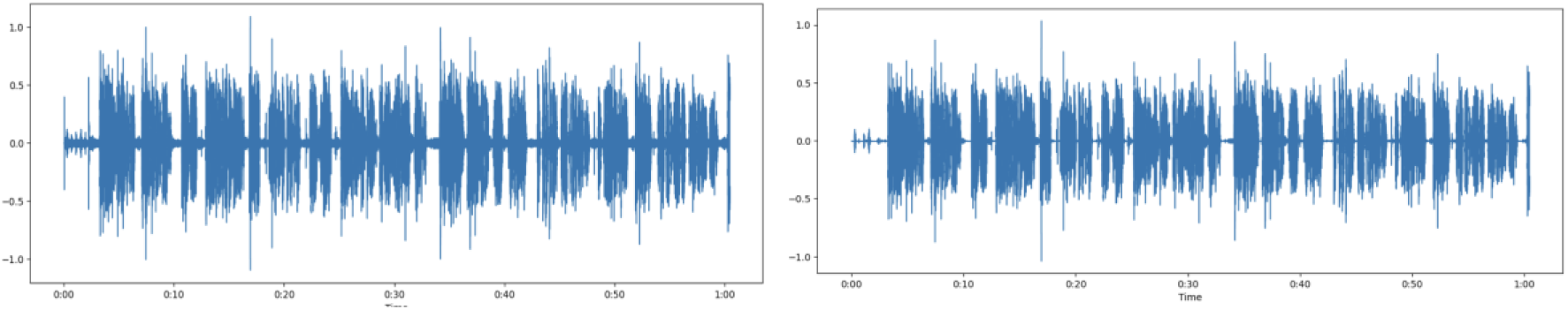
Sound signals before and after the recorded sound is denoised. **(A)** represents the original sound signal which is to be processed while **(B)** is the denoised sound signal which has resulted after undergone the denoiser algorithm.

We then plotted the MFCC (Figure 4) of the denoised signal to be processed for extracting the speech and breath sounds. The waves in the figure represent the speech sounds that are to be demarcated from this signal.

**Fig 4.**
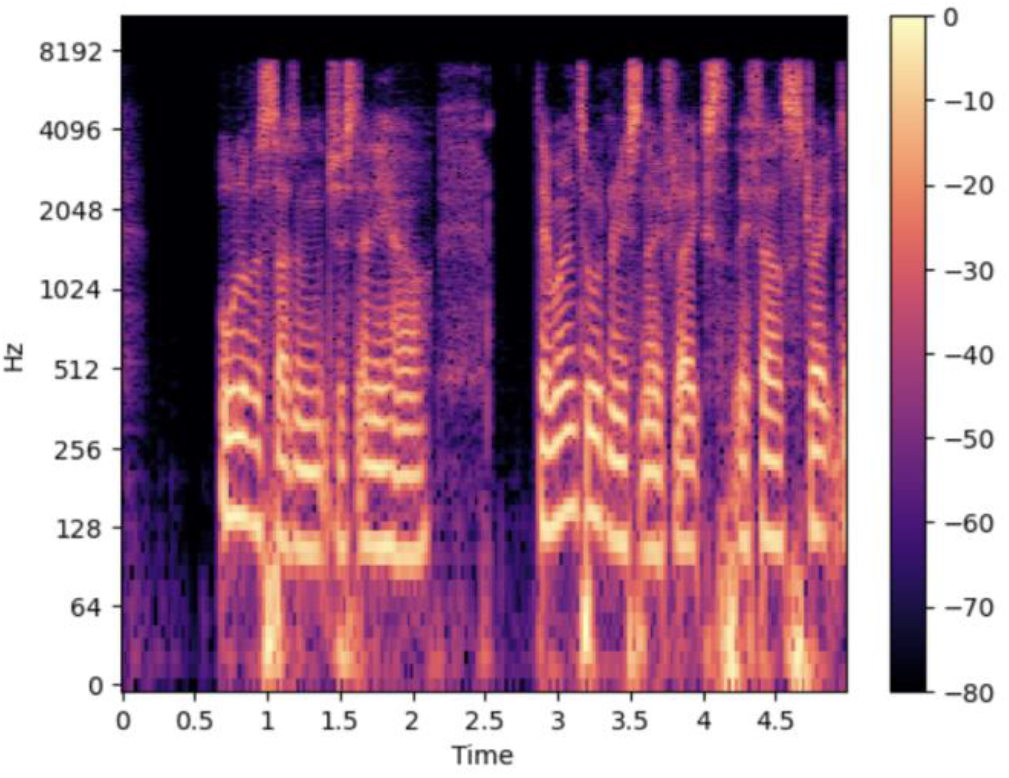
MFCC of the denoised sound file for 5 seconds on a hot scale, the lighter tone on the scale dominantly between 0 Hz - 500 Hz are the speech sounds in the denoised signal while the other darker shades are the breath sounds. Speech sounds can be seen at 1 second, 1.5 second, 3.25 second, 4.2 second and 4.6 second within the 5 second clip of the denoised file.

After the signal was processed through the filters and the masks, the resultant foreground and background signals are displayed in Figure 5 and Figure 6 respectively. The foreground signal is the subsequent speech sound that has been filtered from the denoised signal. The remainder of the full signal after subtraction of the speech or foreground signal is the background signal. The background signal consists of the breath sounds filtered from the original recording.

**Fig 5.**
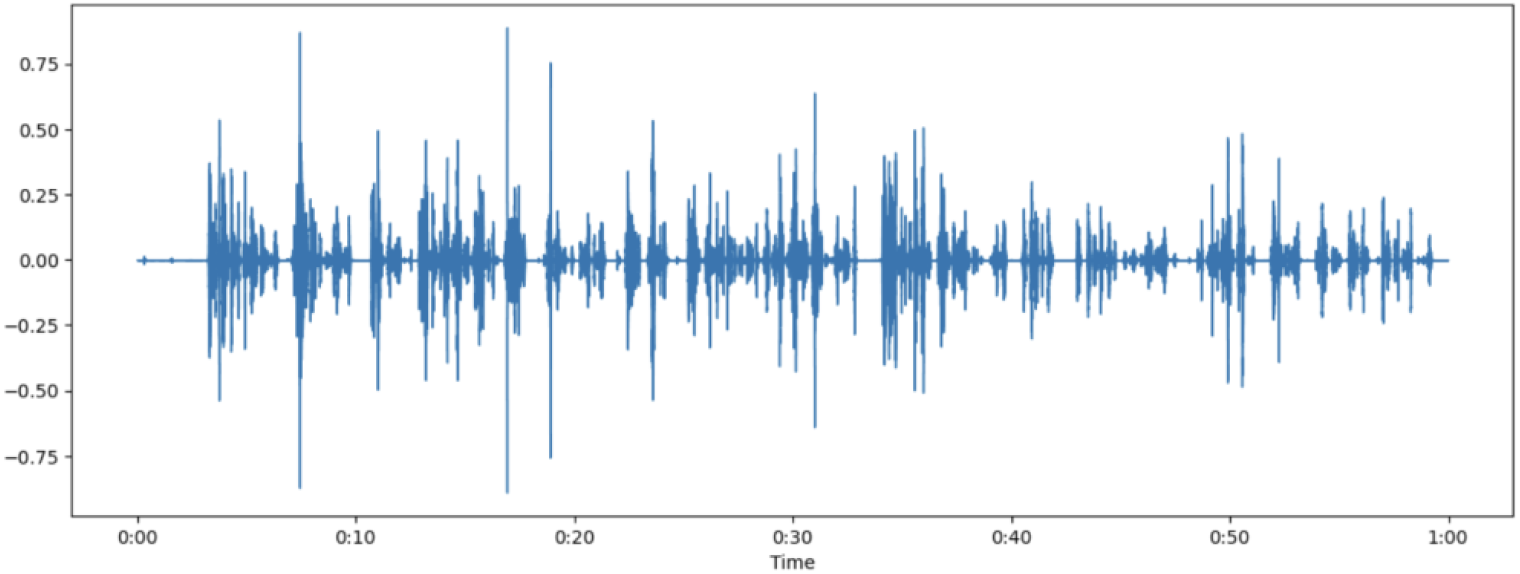
Foreground Signal that represents the speech signal which is filtered from the denoised signal. A combination of STFT and Softmask Weiner filters are used on the original signal to result into the foreground signal.

**Fig 6.**
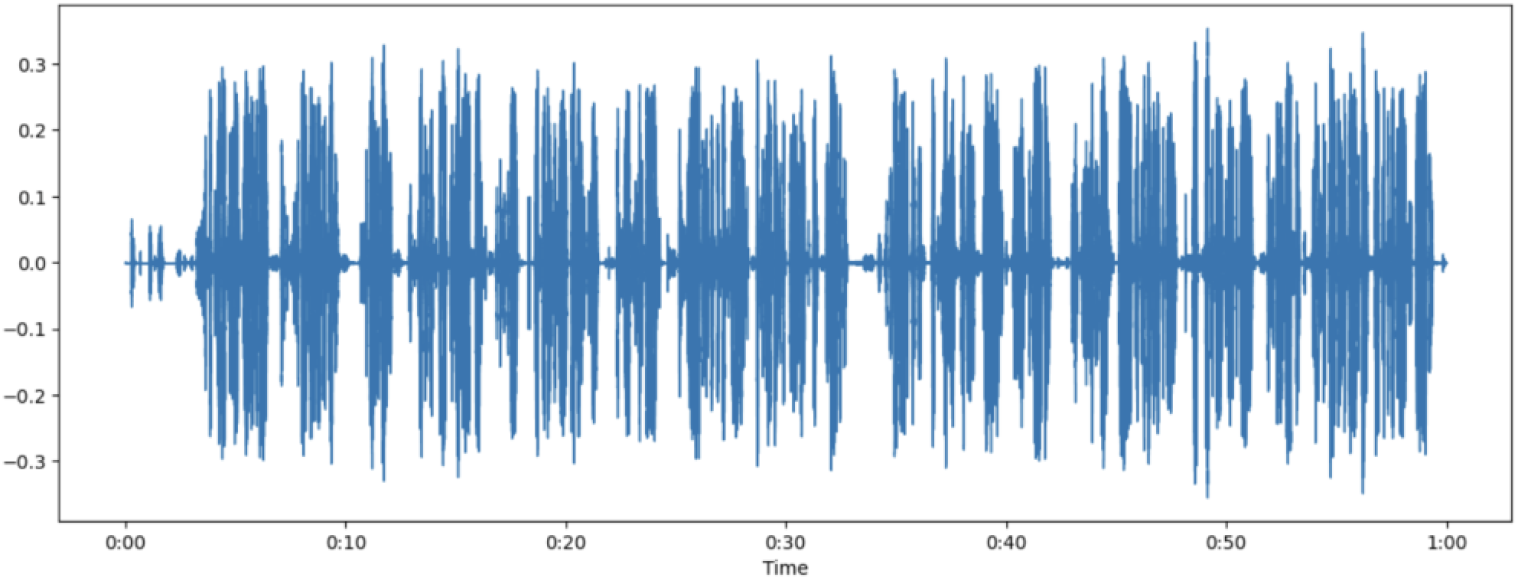
Background Signal that represents the speech signal which is filtered from the denoised signal. A combination of STFT and Softmask Weiner filters are used on the original signal to result into the background signal.

To understand the difference in the MFCC, Figure 7 shows a conjugated signal of the original signal that is labelled as Full Spectrum. While the breath sounds are seen in the Background signal. The Foreground shows the speech sounds filtered from the algorithm.

**Fig 7.**
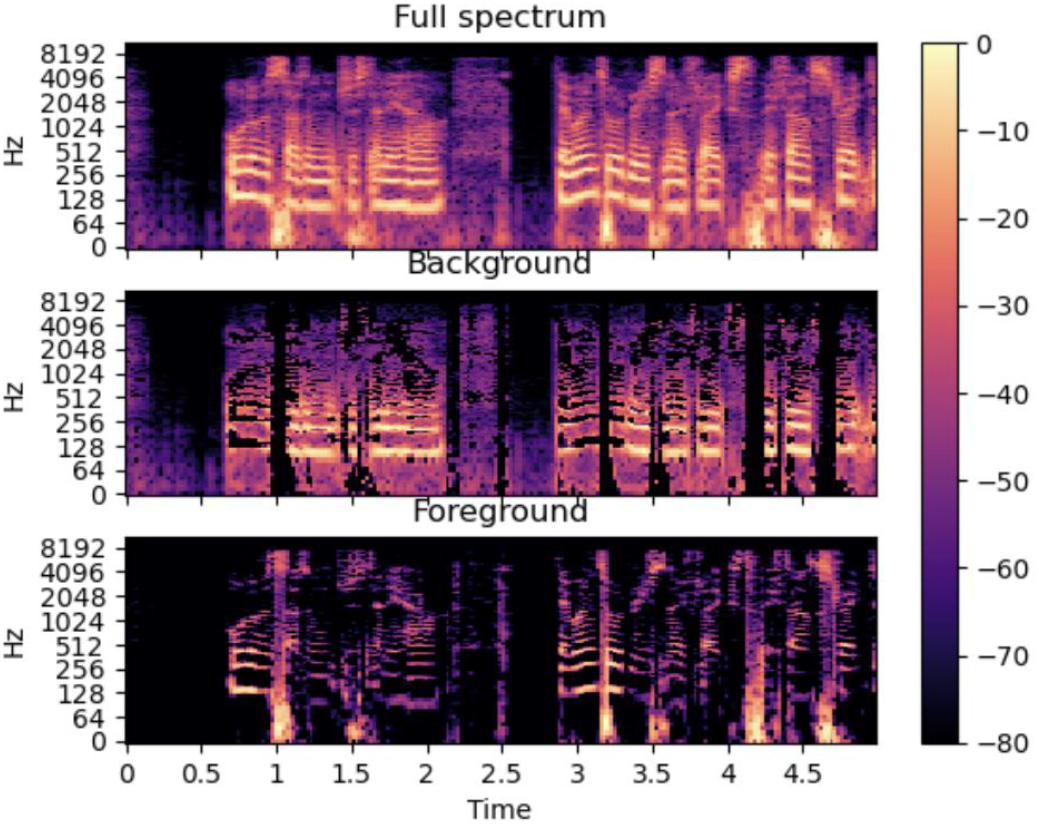
MFCC of the denoised sound (Full Spectrum), Breath Sound (Background) and Speech Sound (Foreground). The displayed images show the difference in signals where the Full Spectrum is the denoised signal, the Background MFCC is the breath sounds where the lighter tones which represent the speech are reduced. The Foreground MFCC is the filtered speech only sounds that have been filtered.

## Discussion

This technique in sound signals along with the denoiser enables to overcome these two drawbacks: First, to make the recording of audio files independent of the use of specialized microphones. The denoiser can process the audio files to filter any noise that would be present in the sound recordings. Secondly, the demarcation algorithm provides a new direction to separate out the speech and breath sounds which was previously done using third-party software such as Audacity and Adobe Audition for processing these files.

The denoiser uses the complex Demucs architecture consisting of encodes, decoders, bidirectional LSTM. This can be used in real-time to denoise any kind of sound signals and can be incorporated with online conferencing, audio communication, signal processing software [14]. One of the objectives was achieved using this architecture.

Filters used in extraction of signals consisted of REPET method for extracting the foreground or the speech signals which consists of Short-time Fourier Transform (STFT) over repeating patterns with a specified window range of ¼. The background signals or the breath sounds were extracted by non-local filtering that was converted to soft mask Weiner filter.

These filters were then multiplied by the denoised signal to output the respective, foreground and background signals. The foreground signals consisted of the speech from the audio file which were perfectly extracted. The background signal consisted of the heavy breath sounds of the patient and the wheezing too is evident.

We experimented with several filters and techniques to achieve the extraction architecture proposed. Speech and breath demarcation is a very prominent area of research especially in disease diagnosis, where symptoms can be gauged in the speech and breath itself. This technique is an attempt to contribute to this area, where till-date the objectives of making the larger models more device independent and remove the dependencies of external software which played the role in signal demarcation. Models can now have this technique implemented to build large-scale diagnosis, prediction and classification algorithms of diseases like bronchitis, asthma, pertussis, COPD and other Upper & Lower Respiratory Tract Infections (URTI and LRTI).

